# Soft tactile stimulation engages parabrachial circuits traditionally associated with aversion

**DOI:** 10.64898/2026.03.15.711870

**Authors:** F Anesten, S Simfors, K Ioneskou, M Hezsö, B Gündogdu, A Tran, A Stjernvall, V Ratiglia, A Almasri, LS Löken

## Abstract

Gentle tactile stimulation is associated with positive affect and social bonding, yet the central circuits engaged by such stimuli remain incompletely understood. The lateral parabrachial nucleus (lePB) is a key hub in ascending affective sensory pathways and is robustly activated by aversive stimuli, including pain. Here, we examined neuronal activation in the lePB and the likewise associated subparafascicular nucleus, parvocellular part (SPFp), following different tactile stimulation paradigms in mice. Behavioral analyses confirmed that the soft touch stimuli used in this study were not aversive: mice displayed low aversive facial grimace scores during brushing and von Frey stimulation compared with noxious heat, and showed a preference for a soft tactile environment in a place preference assay. Neuronal activation was assessed using Fos immunohistochemistry following exposure to brushing-based soft touch, a fur-roll paradigm, innocuous punctate touch (von Frey), or noxious heat. Soft touch protocols robustly increased Fos expression in the lePB compared with home cage controls, whereas innocuous punctate touch did not. Notably, the magnitude of activation produced by brushing-based stimuli was comparable to that induced by noxious heat. Using *Calca*^Cre^ mice, we further found that soft touch recruited a subset of CGRP-expressing neurons in the lePB. In contrast, tactile stimulation produced only modest activation in the SPFp and did not strongly increase overall Fos expression in this region. Together, these findings demonstrate that affective tactile stimulation can engage neuronal populations within ascending parabrachial circuits, including CGRP neurons traditionally associated with nociceptive processing, suggesting that these pathways may encode the salience or affective significance of somatosensory stimuli rather than exclusively aversive input.

## Introduction

Gentle tactile stimulation, such as slow stroking of the skin, is associated with positive affect, social bonding, and stress reduction (Packheiser, Hartmann et al. 2024). In humans, these sensations are mediated by C-tactile (CT) afferents (Loken, Wessberg et al. 2009), a class of unmyelinated low-threshold mechanoreceptors tuned to slow, gentle stroking (Vallbo, Olausson et al. 1993). CT afferents project to brain regions involved in affective and interoceptive processing, including the posterior insular cortex (Olausson, Lamarre et al. 2002, Bjornsdotter, Loken et al. 2009, Morrison, Loken and Olausson 2010). In rodents, the functional analogue of CT afferents are C-low-threshold mechanoreceptors (CLTMRs), which innervate hairy skin and respond selectively to gentle dynamic touch. At the spinal level, CLTMR input is integrated through dorsal horn interneuron networks and can access ascending pathways within the anterolateral system (Choi, Hachisuka et al. 2020, Liu, Qiao et al. 2022). This organization suggests that signals associated with affective touch may converge with nociceptive pathways at early stages of somatosensory processing.

The parabrachial nucleus (PB) is a key node for the processing of sensory signals with affective and motivational value. In rodents, neurons in the lateral external parabrachial nucleus (lePB) and the parvocellular subparafascicular nucleus (SPFp) receive ascending input from spinal and trigeminal pathways. Their projections reach forebrain structures involved in emotional learning, threat detection, and autonomic regulation, including the amygdala, hypothalamus, and bed nucleus of the stria terminalis (Bernard and Besson 1990, Cameron, Polgár et al. 2015) (Kang, Liu et al. 2022, Kang, Liu et al. 2025).

Within these structures, neurons expressing calcitonin gene-related peptide (CGRP) have been strongly implicated in aversive processing and defensive behaviors (Chiang, Bowen et al. 2019, Pauli, Chen et al. 2022). Consistent with this role, a wide range of noxious and aversive stimuli robustly activate neurons in the lePB and SPFp (Palmiter 2018, Condon, Yu et al. 2024, Palmiter 2024).

Recent studies suggest that these nuclei may encode generalized sensory salience rather than modality-specific nociceptive signals, as diverse aversive stimuli activate overlapping neuronal populations within these regions (Kang, Liu et al. 2022, Kang, Liu et al. 2025). This raises an important question: do these circuits exclusively signal aversion, or can they also be engaged by non-noxious tactile stimuli with positive affective significance?

In rodents, soft tactile stimuli such as brushing of the fur or contact with soft materials evoke behaviors consistent with positive affect (Choi, Hachisuka et al. 2020, Liu, Qiao et al. 2022). Here, we investigated how different forms of soft touch influence neuronal activation in the lePB and SPFp, and whether this activation involves CGRP-expressing neurons.

Using Fos immunohistochemistry, we compared multiple soft touch protocols with innocuous punctate touch and noxious heat. To assess the affective valence of these stimuli, we complemented neural measurements with behavioral assays including place preference and facial grimace scoring. Our results show that soft touch robustly activates parabrachial neurons, and to some extent also activates the CGRP-expressing population within the lePB, despite lacking behavioral signatures of aversion. These findings suggest that parabrachial circuits traditionally associated with negative valence may also participate in processing affectively salient, non-aversive tactile signals.

## Methods

### Animals

Male and female heterozygous *Calca*^Cre^ mice (8–12 weeks old; JAX #033168) mice express nuclear-localized cre recombinase:EGFP fusion protein targeted to mouse *Calca* gene (here referred to as CGRP-GFP) and FosTRAP2 mice (JAX #030323) on a C57BL/6J background were bred in-house at the Facility for Experimental Biomedicine, Sahlgrenska Academy, University of Gothenburg. FosTRAP mice were used without tamoxifen administration and therefore functioned as wild-type mice.

Animals were group-housed under standardized conditions (12 h light/dark cycle; lights on at 07:00; temperature 21 °C; humidity 50–60%) with ad libitum access to standard chow and water.

All experimental procedures were approved by the local Animal Ethics Committee at the University of Gothenburg (ethical approval number 4534/22) and were conducted in accordance with national and European Union guidelines for the care and use of laboratory animals.

### Behavior Protocols

All behavior protocols were performed between 9-11 am in the same room to which the mice had previously been habituated over a course of 4-5 days. Habituation consisted of remaining in the home cage in the test room for 15 mins with a permeable lid, for the mice to get used to the smells and sounds of the room as well as the experimenter. After this habituation, mice were transferred to the experimental cage (see each test for specifications) and allowed to acclimatize for 15 mins. Mice who were exposed only to sitting in their home cage were used as a control group.

### Brush

The hairy back skin was gently stroked with a hand-held soft brush, moving from the nape of the neck to the lumbar enlargement region at constant speed (18-22 cm/s) and force (maximum 23-25 mN) as described by Liu et al (ref). Three bouts of 90 s of brushing, with a 300 s rest period in between each bout, were performed. The first of these bouts was filmed with a Sony Handycam CX405. Each mouse had its own brush to avoid odor contamination. The protocol was repeated for a period of ten days over the span of two weeks. On the tenth day mice were perfused and sacrificed 90 min after the protocol was completed.

### Brush + Non-Noxious Heat and Blanket

Mice were placed on a hot plate set to 37°C. A soft blanket of similar coloration and texture to the fur of a C57BL/6J mouse was placed on the hot plate, covering roughly half of it. Brushing was performed as described above. Each mouse had its own brush and blanket to avoid odor contamination. The protocol was repeated for a period of ten days over the span of two weeks. On the tenth day mice were perfused and sacrificed 90 min after the protocol was completed.

### Blanket Cage

Mice were placed in a plastic cage that was divided into two equal sides with a separator. Individual cages were used for each mouse to avoid odor cross-contamination. One side of the cage was covered in a soft blanket as described above. The other side was left bare but with the transparent cage bottom sitting atop the same kind of blanket. For habituation, the mouse was placed alternatingly on the blanket side and the bare side, 15 min for each side. This protocol was repeated daily over a period of five days. On the fifth day the separator was removed, allowing the mouse to freely choose which side to dwell in for 15 min. The test on the fifth day was recorded as described above, and time spent by the mouse on either the blanket or the bare area was then measured.

### Fur Roll

Mice were allowed free access to a cardboard cylinder clad in a blanket (as above) placed inside a blanket cage (as above) for a duration of 30 min. Each mouse had its own cage and roll to avoid odor contamination. This protocol was repeated daily over a period of five days. On the fifth day, mice were first left inside the blanket cage with the blanket-clad roll for 15 min. The latter 15 min of the test were recorded as described above. Mice were perfused and sacrificed 90 min after the end of the experiment.

### Noxious Heat (Hot Plate)

Mice were placed on a hot plate covered by a sturdy piece of cardboard inside a plastic cylinder. Mice were left to acclimatize for 15 min, and thereafter the temperature was slowly ramped up to 50°C over approximately 5 minutes. The cardboard plate was removed as the temperature reached close to target and the mouse left on the bare hot plate for up to maximum 15 s or until pain behavior was observed (jumping, hindpaw lifting, paw licking). The cardboard plate was then reintroduced and the mouse allowed 90 s of recovery time. This was repeated seven to eight times. Mice were perfused and sacrificed 90 min after the end of the experiment.

### von Frey

Mice were acclimatized to the testing room and the raised metal grid where they were placed for the experiment over a period of four days. On the fifth day, mice were left in their home cage in the testing room for 15 min after which they were transferred to plastic cylinders atop the raised metal grid and left to acclimatize for 15 min. The right hind paw was stimulated using a 0.4 g von Frey filament, enough to be felt by the mouse but not be painful (Deuis, Dvorakova and Vetter 2017). Stimulation went on until the mouse exhibited paw withdrawal, and repeated over a period of 15 min interspersed with resting periods. Mice were perfused and sacrificed 90 min after the end of the experiment.

### Tissue preparation and Immunohistochemistry

Mice were deeply anaesthetized with a mixture of domitor and ketamine, and perfused transcardially with 10 mL PBS, followed by 30 mL 4% paraformaldehyde. The brain was removed and post fixed in 4% paraformaldehyde for 3 h. Tissue was then transferred to a 30% sucrose solution in PBS until sectioning. Coronal 30 μm thick serial sections of the midbrain and thalamus were cut using a Leica CM3050S cryostat (Leica CM3050 S, Leica Biosystems, Wetzlar, Germany) and stored in PBS.

Sections were mounted on SuperFrost slides and blocked for 1.5 h with 10% normal goat serum (abcam, Cambridge, UK) in PBS with 0.3% Triton-X-100 (Perkin Elmer, Waltham, MA, USA). Sections were incubated with primary antibodies (Rabbit anti-cfos 1:2000, RRIID:AB_2247211; Chicken anti-GFP 1:3000, RRID:AB_300798) overnight at room temperature. After rinsing with PBS, sections were incubated for 2 h with secondary antibodies (Goat Anti-chicken Alexa Fluor 488 1:1000 and Goat Anti-rabbit Alexa Fluor 555 1:1000, A-10680, ThermoScientific, Waltham, MA, USA) with 1% normal goat serum in PBS with 0.3% Triton-X-100. Sections were rinsed with PBS, mounted with Fluoromount G and coverslipped.

### Microscopy and Cell Counting

Sections were imaged using a Nikon Eclipse Ti2 inverted microscope with NIS-Elements software using a 10x or 20x lens, using the same settings for exposure time and laser intensity for all images. Images were adjusted for brightness and contrast in FIJI (version 2.1.0). Representative images from each of the two nuclei were obtained, one from each mouse for each distance from bregma. The appropriate extention of the nucleus was manually delineated and the cells were then counted cells manually. A researcher blind to the behavior protocol then recounted a sample of slides from each group to ensure unbiased counting. Only cells with a light intensity between 130-300 that had a uniform shape were counted.

### PainFace Analysis

Mice were filmed during behavior protocols as described above. Movies were cut using VLC Media Player. The resulting clips had the mouse facing the camera during the behavioral stimulus with its face in clear view. Video clips were then stitched together using QuickTime Player and uploaded to the PainFace website (McCoy, Park et al. 2024). For analysis using the PainFace algorithm the following settings were used: fau model ID default(20250513-general), pain-mgs model ID default(20221115-black) and Sample Rate High (1 Frame/Second). Scores (0-2) from the four face parts analyzed by the algorithm (Orbital, Nose, Ears, Whiskers) were extracted from the completed analyses and imported into Excel. After removing frames where the face area was not visible (defined as a score of −1), average scores were obtained from each mouse and behavior test for each face area, as well as a total score for all four face areas.

### Statistics

All statistical analyses were conducted using IBM SPSS Statistics (Version 30.0.0.0, IBM Corp., Armonk, NY, USA). Data were first tested for normality using the Shapiro–Wilk test and for homogeneity of variances using Levene’s test. One-way ANOVA was performed to examine the effect of treatment (Control, Brush, Brush+, Fur roll, Noxious heat, and von Frey) on Fos expression, as well as proportional overlap between Fos and CGRP, and vice versa. Tukey’s HSD post hoc tests were used for pairwise group comparisons when the assumption of equal variances was met. When the assumption of homogeneity of variance was violated, Welch’s ANOVA was used instead, followed by Games–Howell post hoc comparisons. Post hoc analyses were conducted for all treatment groups. All results are presented as mean ± SEM, with statistical significance set at p < 0.05. Graphs were generated using Prism (version 10.6.1).

## Results

### Soft touch stimuli are not aversive to mice

Because the neural responses reported below compare activity evoked by soft touch and noxious heat, we first sought to determine whether the soft touch stimulus might be perceived as aversive. As unpleasant stimuli in mice are known to increase facial grimacing, we quantified grimace scores during the different behavioral protocols.

Mice were filmed during the Brush, Noxious Heat, and von Frey protocols. As a baseline, mice were filmed during unstimulated conditions in the test cage used for the Brush and von Frey assays (see figure 1A). Because the stimulation periods were brief (~15–90 s), frames were extracted during ongoing stimulation when the mouse was facing the camera. This resulted in relatively few frames in which all four facial action units were clearly visible; therefore, partial frames in which one to three facial regions were visible were also included. Scores for each individual facial action unit were summed to obtain an average grimace score.

**Figure 1.**
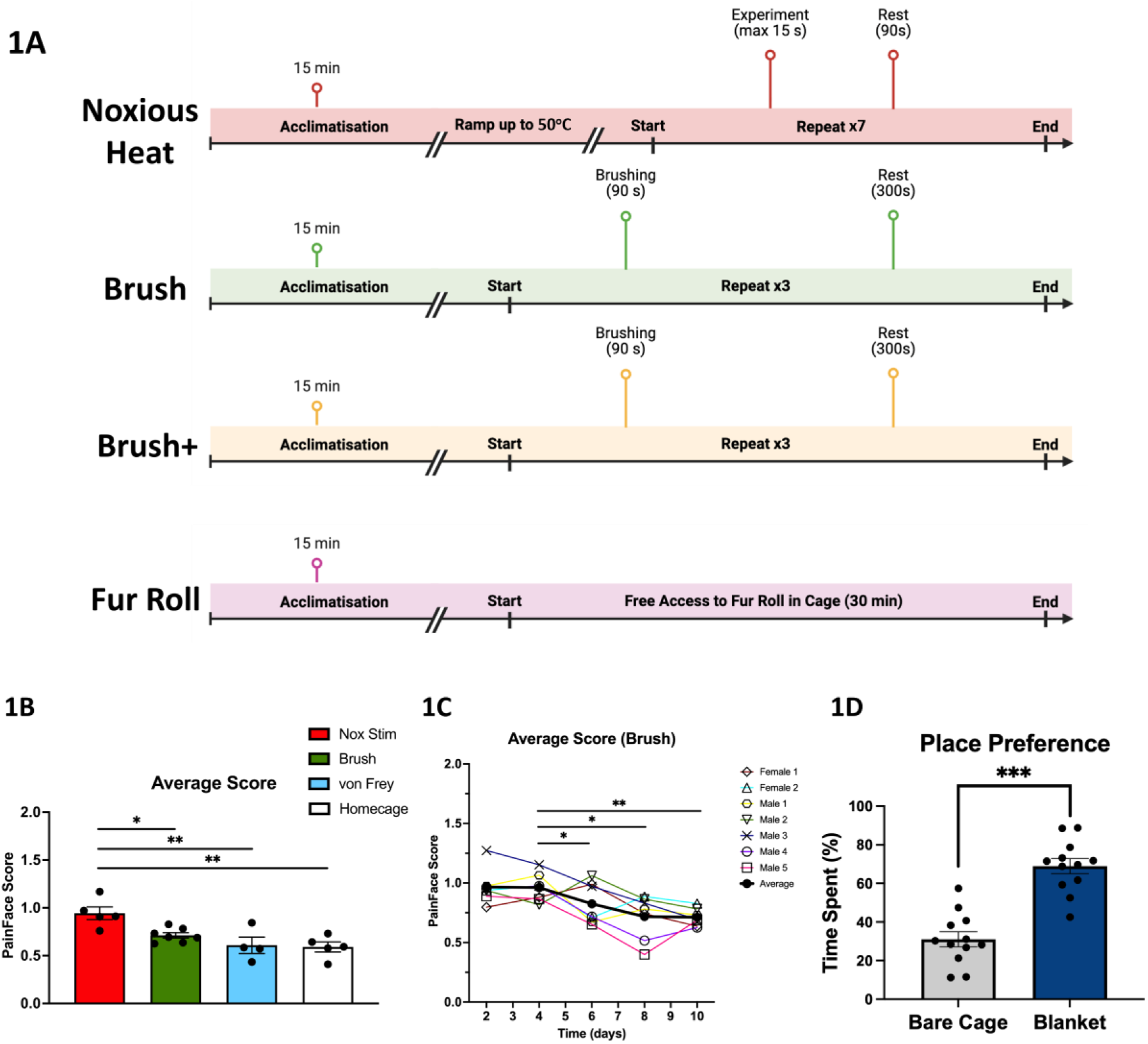
Soft touch stimuli are not aversive to mice. **(A)** Timelines for the behavior protocols from top to bottom: Noxious Heat, Brush, Brush +, Fur Roll. **(B)** Average grimace scores of mice exposed to noxious heat are significantly higher in mice exposed to noxious heat (n=5, M=0.94, SE=0.07) as compared with Brush (n=7, M=0.71, SE=0.03, *p* = 0.026), von Frey (n=4, M=0.61, SE=0.09, *p* = 0.005), or Baseline (n=5, M=0.59, SE=0.05, *p* = 0.002) conditions. **(C)** Mice (n=7) exposed to the Brush protocol display lower average grimace scores over time. Mice displayed higher average grimace scores on day 2 (M=0.97, SE=0.06) compared with day 8 (M=0.72, SE=0.07, *p* = 0.026) and day 10 (M=0.71, SE=0.03, *p* = 0.022), and on day 4 (M=0.96, SE=0.04) compared with day 8 (*p* = 0.029) and day 10 (values, *p* = 0.025). **(D)** In a conditioned place preference test, mice spent significantly more time on the blanket-covered side than on the bare floor (n= 12. Bare side M=31, SE=3.91, Blanket side M=69, SE=3.91. *p* < 0.001). * = *p* < 0,05, ** = *p* < .005, *** = *p* < .001.

When comparing average grimace scores across conditions using a one-way ANOVA, mice exposed to noxious heat exhibited significantly higher grimace scores than mice in the Brush (*p* = 0.026), von Frey (*p* = 0.005), or Baseline (*p* = 0.002) conditions (figure 1B).

We next examined whether grimace scores during the Brush protocol changed over time, as we hypothesized that the unfamiliarity of the stimulus might initially increase grimacing in the early days of habituation. Grimace scores were extracted from days 2, 4, 6, 8, and 10 of Brush protocol habituation. We found a significant difference between early and late days of habituation, such that mice displayed higher average grimace scores on day 2 compared with day 8 (*p* = 0.026) and day 10 (*p* = 0.022), and on day 4 compared with day 8 (*p* = 0.029) and day 10 (*p* = 0.025) (figure 1C). No other comparisons were significant.

Because facial recordings could not be obtained during the fur-roll condition, we next assessed whether mice showed avoidance or preference for a soft tactile environment using a place preference assay. Mice were habituated as described in Methods. On the test day, mice spent significantly more time on the blanket-covered side than on the bare floor (69% on the blanket side and 31% on the bare side (*p* < 0.001, figure 1D)).

Taken together with previously published findings that investigate the same kind of protocols, we conclude that the soft touch protocols are non-aversive.

### Soft touch engages neurons in the lateral external parabrachial nucleus

The lateral external parabrachial nucleus (lePB) (figure 2A) is an early hub in ascending affective sensory pathways and is robustly activated by aversive stimuli (Palmiter 2018, Condon, Yu et al. 2024, Palmiter 2024). We therefore asked whether soft touch stimuli would also engage neurons in this region.

**Figure 2.**
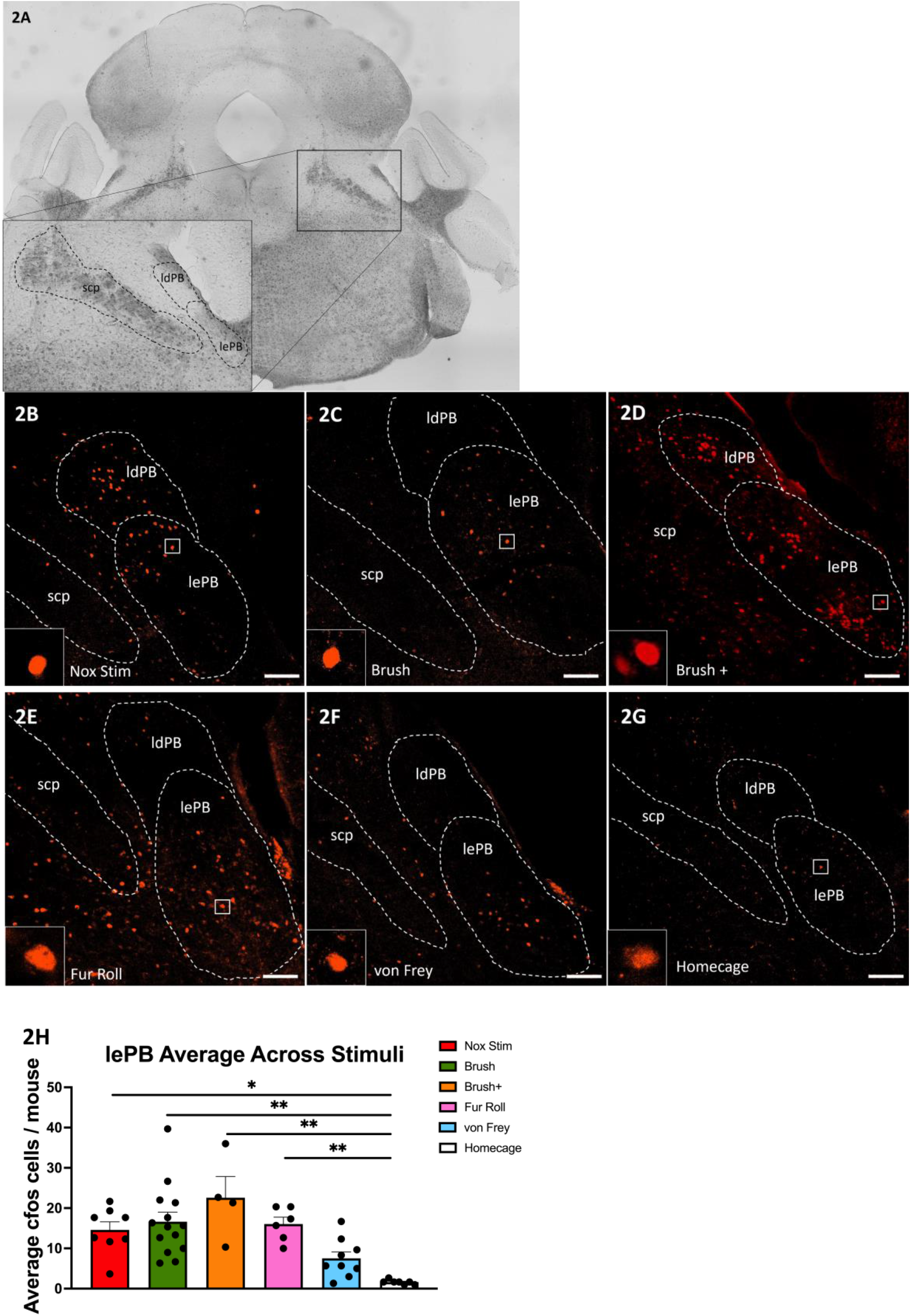
Soft touch engages neurons in the lateral external parabrachial nucleus. **(A)** Anatomical overview of the mouse lateral parabrachial nucleus, showing its dorsal (ldPB) and external (lePB) subdivisions, as well as the superior cerebellar peduncle (scp). **(B-G)** Representative micrographs showing Fos expression (red) after **(B)** noxious heat, **(C)** Brush, **(D)** Brush +, **(E)** blanket roll, **(F)** von Frey or **(G)** homecage conditions (scalebars 100μm). **(H)** Noxious heat (n=8, M=43.75, SE=6.06, *p* = 0.004), Brush (n=14, M=49.86, SE=7.10, *p* < 0.001), Brush + (n=4, M=67.75, SE=15.77, *p* < 0.001), Fur Roll (n=6, M=48.17, SE=5.10, *p* = 0.003) all show significantly increased Fos expression as compared with homecage (n=7, M=4.86, SE=0.63). In contrast, low-force von Frey stimulation did not increase Fos expression (n=9, M=22.56, SE=4.80, p=0.5). * = *p* < 0,05, ** = *p* < .005.

To test this, neuronal activation was assessed using immunohistochemistry for the immediate early gene Fos following exposure to our battery of tactile stimulation protocols. Mice were exposed to one of three soft touch conditions: brushing of the fur (Brush), brushing combined with mild non-noxious warmth and a blanket (37 °C, Brush +), and a Fur Roll condition in which animals were allowed to enter a fur-clad roll. As a comparison condition representing innocuous non-affective touch, another cohort of mice were stimulated with a low-force von Frey filament (0.4 g). Another cohort of mice were exposed to a noxious heat condition to serve as a positive control. Fos–positive neurons were quantified at three rostrocaudal levels of the lePB (−5.0, −5.1, and −5.2 mm from bregma), and mean counts across levels were used for statistical comparisons.

One-way ANOVA revealed a significant effect of condition (*F* (5, 42) = 8.17, *p* < 0.001). Post hoc Tukey tests showed that all soft touch protocols and noxious heat significantly increased the number of cfos–positive neurons in the lePB compared with home cage controls: Brush (*p* < 0.001), Brush + (*p* < 0.001), Fur Roll (*p* = 0.003) Noxious heat (*p* = 0.004). In contrast, low-force von Frey stimulation did not increase Fos expression relative to home cage mice (*p* = 0.5) (figure 2 B-H).

The post-hoc comparison of the stimulation protocols further revealed that brushing-based soft touch produced significantly more Fos expression compared to von Frey stimulation (Brush, *p* = 0.029; Brush +, *p* = 0.006), whereas Fur Roll did not differ significantly from von Frey (*p* = 0.166). Consistent with previous reports, noxious heat significantly increased Fos expression relative to home cage controls (*p* = 0.039). Notably, Fos expression evoked by noxious heat did not differ from those induced by the soft touch protocols (Brush *p* = 0.982; Brush + Heat, *p* = 0.378; Fur Roll, *p* = 0.998).

Taken together, these findings confirm that noxious heat activates the lePB and demonstrate that soft touch, particularly brushing-based stimulation, also robustly activates lePB neurons, exceeding the activation produced by innocuous punctate touch.

### Soft touch activates a subset of CGRP neurons in the lateral external parabrachial nucleus

To determine whether neurons activated by soft touch included CGRP-expressing neurons, two of the soft touch protocols (Brush and Fur Roll) were repeated in CalcaCre mice. This allowed us to assess the overlap between Fos–positive neurons and CGRP-expressing neurons in the lePB.

We first quantified the proportion of Fos–positive neurons that also expressed CGRP. A one-way ANOVA revealed a significant effect of stimulation condition on this proportion (*F* (4, 21) = 4.9, *p* = 0.006). Post hoc Tukey tests showed that the Fur Roll and noxious heat protocols significantly increased the proportion of Fos-positive neurons that were CGRP-positive compared with home cage controls (*p* = 0.008 and *p* = 0.004, respectively), whereas Brush showed a similar trend (*p* = 0.055). Although von Frey stimulation produced fewer Fos-positive neurons overall, the proportion of Fos–CGRP co-localization did not differ significantly from that observed in the Brush, Fur Roll, noxious heat, or home cage conditions.

Next, we examined the proportion of CGRP neurons that were Fos-positive following stimulation. Stimulation condition again significantly affected this measure (one-way ANOVA, *F* (4, 21) = 4.8, *p* = 0.007). Post hoc comparisons revealed that both Brush and Fur Roll significantly increased the proportion of CGRP neurons expressing Fos compared with home cage controls (Brush, *p* = 0.014; Fur Roll, *p* = 0.011) (figure 3 A-G).

**Figure 3.**
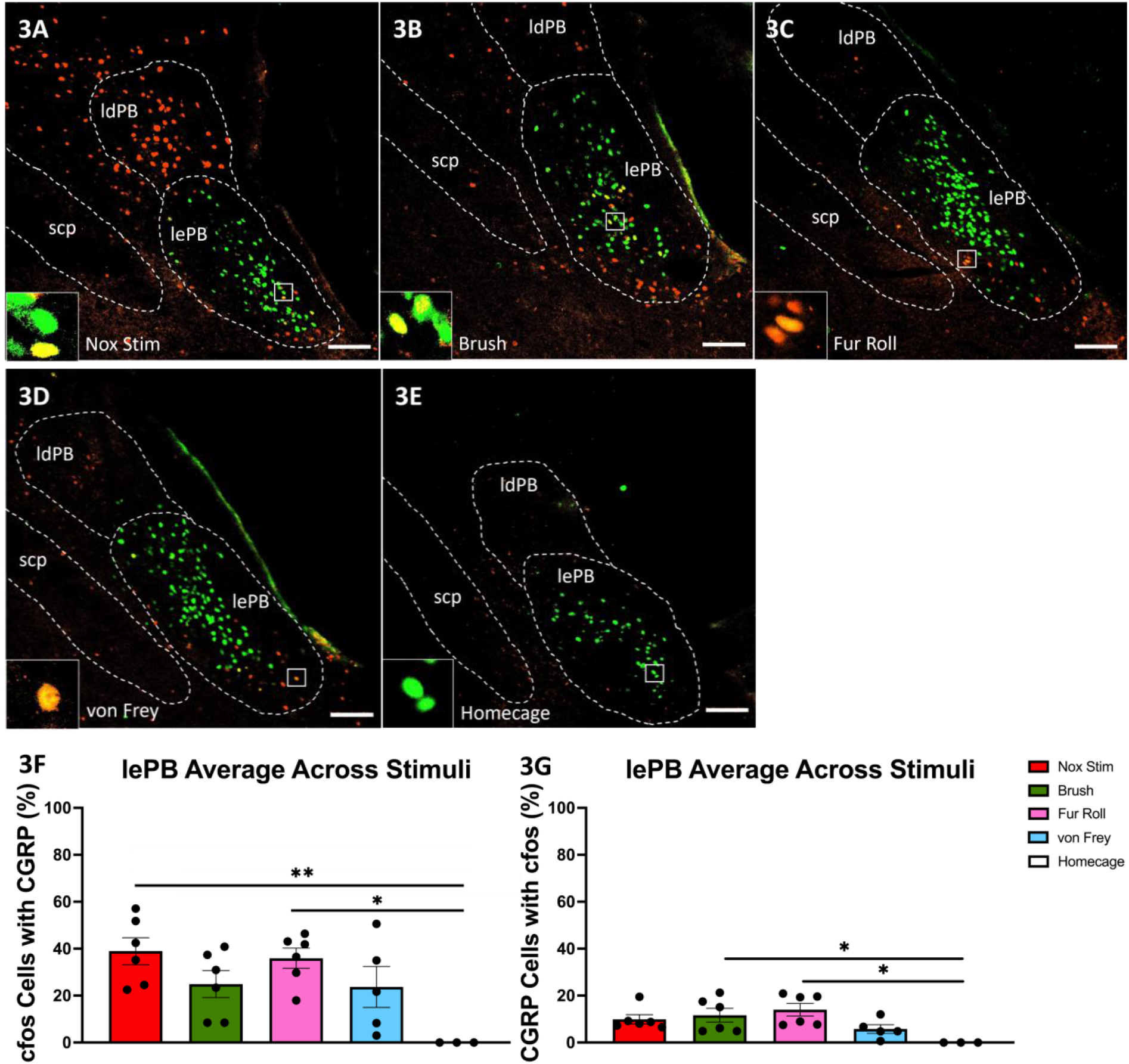
Soft touch activates a subset of CGRP neurons in the lateral external parabrachial nucleus. **(A-E)** Representative micrographs showing Fos expression (red) and CGRP-GFP (green) after **(A)** noxious heat, **(B)** Brush, **(C)** blanket roll, **(D)** von Frey or **(E)** homecage conditions (scalebars 100μm). **(F)** Noxious heat (n=6, M=38.94, SE=5.77, p=0.004), and blanket roll (n=6, M=35.95, SE=4.31, p=0.008) show a significantly increased proportion of Fos-expressing neurons that also express CGRP, as compared with homecage (n=3, M=0, SE=0). No such difference was found for Brush (n=6, M=27.71, SE=4.72) or von Frey (n=5, M=23.72, SE=8.71). **(G)** Brush (M=13.63, SE=2.69, p=0.014) and blanket roll (M=14.01, SE=2.67, p=0.011) show a significantly increased proportion of CGRP-expressing neurons that also express Fos, as compared with homecage (M=0, SE=0). No such difference was found for noxious heat (ME=9.91, SE=1.96) or von Frey (M=5.79, SE=1.85). * = *p* < 0,05, ** = *p* < .005.

To determine whether this recruitment was specific to affective touch, we compared these results with the von Frey protocol, a measure of innocuous punctate touch. In contrast to Brush and Fur Roll, neither von Frey nor noxious heat significantly increased the proportion of CGRP neurons that were Fos-positive compared with home cage controls, and these proportions did not differ significantly from those observed in the other conditions. Together, these results indicate that soft touch stimuli recruit a subset of CGRP neurons in the lePB.

### Parvocellular subparafascicular nucleus neurons are activated by noxious heat but modestly by tactile stimulation

The SPFp (figure 4A) also contains a population of CGRP-expressing neurons and receives sensory inputs similar to those of the lePB (Kang, Liu et al. 2022, Kang, Liu et al. 2025). We therefore performed the same analyses in the SPFp using tissue from CalcaCre mice.

**Figure 4.**
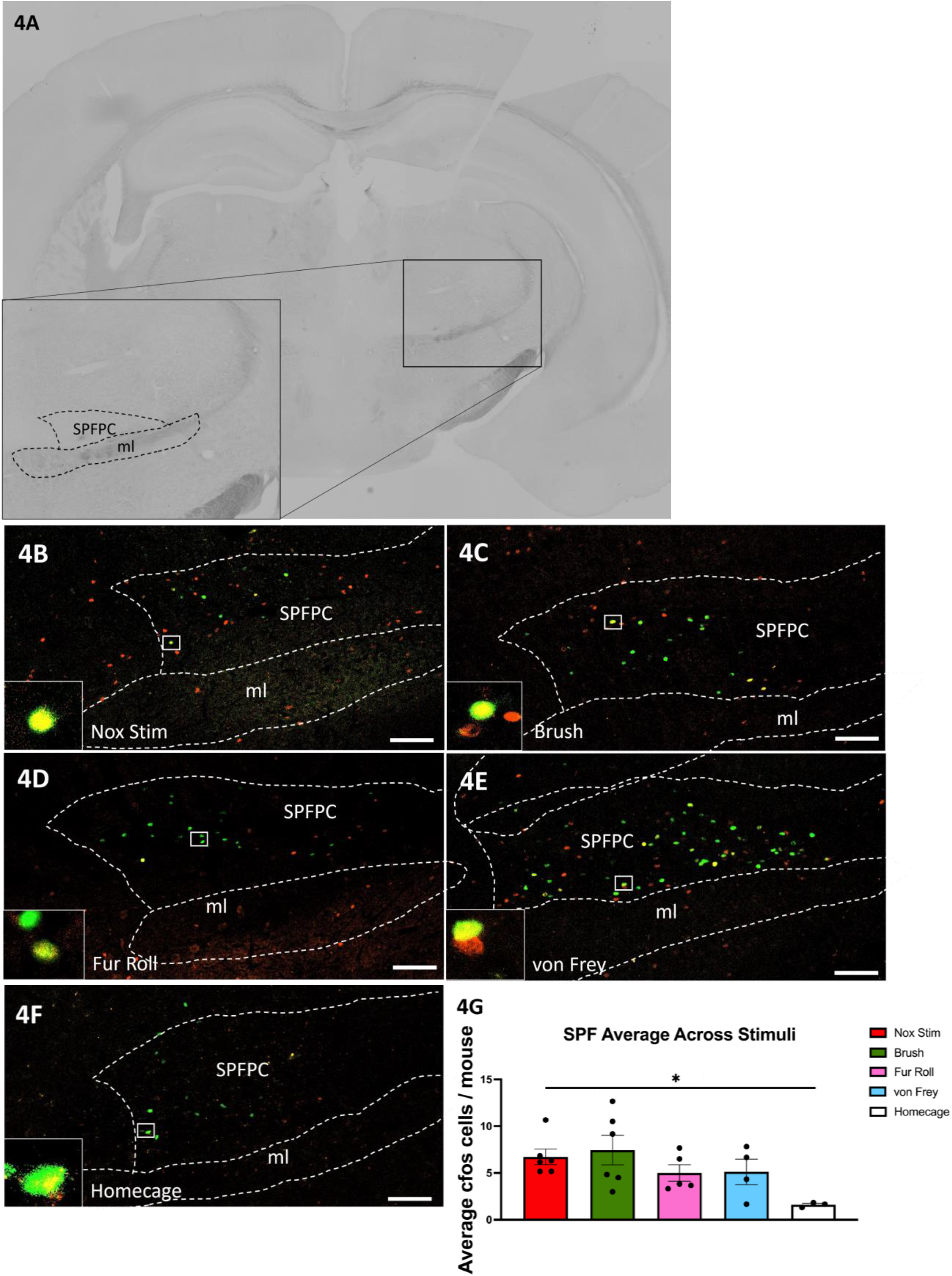
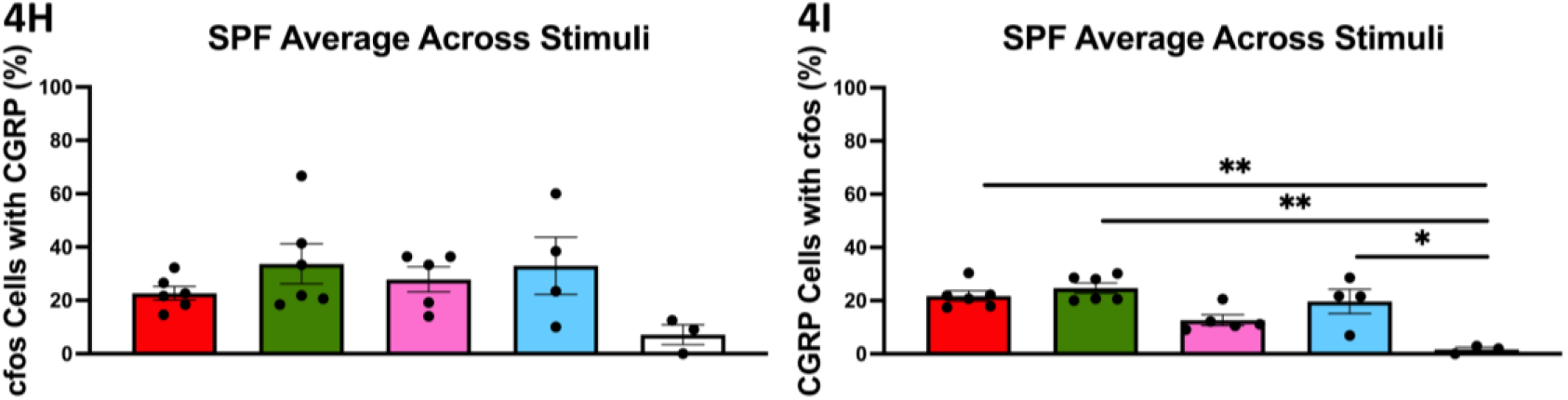
Parvocellular subparafascicular nucleus neurons are activated by noxious heat but modestly by tactile stimulation. **(A)** Anatomical overview of the mouse parvocellular subparafascicular nucleus (SPFPC) and the medial lemniscus (ml). **(B-F)** Representative micrographs showing Fos expression (red) and CGRP-GFP (green) after **(A)** noxious heat, **(B)** Brush, **(C)** blanket roll, **(D)** von Frey or **(E)** homecage conditions (scalebars 100μm). **(G)** Noxious heat (n=6, M=6.72, SE=0.83, p=0.008) shows significantly increased Fos expression as compared with homecage (n=3, M=1.61, SE=0.15). Brush (n=6, M=7.44, SE=1.58), blanket roll (n=5, M=5.00, SE=0.87) and von Frey (n=4, M=5.13, SE=1.36) showed no such differences. **(H)** No difference in the proportion of Fos-expressing neurons that also expressed CGRP was found in any of the conditions, as compared to homecage. **(I)** Noxious heat (M=22.44, SE=2.90, p=0.003) and Brush (M=22.62, SE=2.53, p=0.003) show a significantly increased proportion of CGRP-expressing neurons that also express Fos, as compared with homecage (M=1.65, SE=0.87. No such difference was found for blanket roll (M=12.64, SE=1.99) or von Frey (M=19.19, SE=5.29). * = *p* < 0,05, ** = *p* < .005.

We first quantified the number of Fos-positive neurons following tactile stimulation. Because homogeneity of variance was violated, Welch’s ANOVA was used. Welch’s ANOVA revealed a significant effect of stimulation condition on the mean number of Fos-positive neurons in the SPFp (Welch’s *F* (4, 8.4) = 13.7, *p* < 0.001) (figure 4G). Games–Howell post hoc comparisons showed that the noxious heat group had significantly higher Fos counts than home cage controls (*p* = 0.008), whereas none of the tactile stimulation conditions differed significantly from controls.

We next examined whether neurons activated by tactile stimulation in the SPFp also expressed CGRP. A one-way ANOVA revealed no significant effect of stimulation condition on the proportion of Fos-positive neurons that co-expressed CGRP (*F* (4, 19) = 1.8, *p* = 0.176).

Finally, we assessed the proportion of CGRP neurons that expressed Fos following stimulation. A one-way ANOVA revealed a significant effect of stimulation condition on this measure (*F* (4, 19) = 6.5, *p* = 0.002) (figure 4H-I). Post hoc comparisons showed that the Brush protocol significantly increased the proportion of CGRP neurons that were Fos-positive compared with home cage controls (*p* = 0.003), whereas Fur Roll did not (*p* = 0.211). Similar increases were observed following von Frey (*p* = 0.022) and noxious heat (*p* = 0.003) stimulation. Together, these results suggest that tactile stimulation recruits a small subset of CGRP neurons in the SPFp without producing a strong increase in overall neuronal activation in this region.

## Discussion

We here show that several forms of tactile stimulation, including soft affective touch, recruit neuronal populations in the lePB and modestly in the SPFp. Importantly, innocuous punctate touch did not significantly increase neuronal activation in the lePB compared with home cage controls, suggesting that soft touch protocols carries an affective component or are more salient stimuli. Consistent with previous studies, noxious heat also increased activation of lePB neurons (Chiang, Nguyen et al. 2020).

Because CGRP neurons in these regions are typically linked to aversive processing, we first considered whether the soft touch stimuli might be perceived as unpleasant by the animals. However, several observations argue against this interpretation. Previous studies have shown that mice prefer both soft brushing and blanket-lined environments over unstimulated conditions (Liu, Qiao et al. 2022, Liu, Rahman et al. 2025). Consistent with these reports, mice in our place preference assay spent significantly more time on the blanket-covered side of the cage than on the bare floor. There has been considerable work done in the field of mouse grimace analysis in the context of stimuli with both positive and negative valence (Langford, Bailey et al. 2010, Dolensek, Gehrlach et al. 2020, Le Moene and Larsson 2023). In our hands, facial grimace analysis revealed no increase in aversive grimacing during brushing compared with baseline or punctate non-affective touch conditions. Grimacing in mice also decreased progressively during habituation to brushing. This potentially indicates a change towards positive valence. Together, these findings suggest that the soft touch protocols used in the present study are unlikely to be aversive.

Previous work has demonstrated that CGRP neurons in both the parabrachial nucleus and the SPFp contribute to the affective and motivational dimensions of pain (Palmiter 2018, Kang, Liu et al. 2025). In the present study, noxious heat, and soft touch produced similar levels of activation within CGRP neuronal populations in the lePB. In contrast, activation within the SPFp was more limited, with overall neuronal activation primarily driven by noxious heat. These findings suggest that although both regions contain CGRP neurons associated with affective processing, the lePB may be more responsive to non-aversive tactile input than the SPFp.

An additional consideration when interpreting Fos expression is that immediate early gene induction often reflects relatively strong or sustained neuronal activation. Classic studies have shown that noxious stimulation reliably produces robust Fos induction throughout pain pathways, whereas weaker or transient sensory inputs may not consistently engage these transcriptional responses (Hunt and Mantyh 2001). In this context, the observation that gentle tactile stimulation was sufficient to induce Fos expression in a subset of CGRP neurons in the lePB suggests that these neurons can be recruited even by relatively mild sensory input. One possible explanation for the recruitment of CGRP neurons by brushing is that this stimulus engages C-low-threshold mechanoreceptors (CLTMRs), unmyelinated afferents tuned to gentle dynamic stimulation of hairy skin. Activation of CLTMR pathways in mice has been shown to promote behavioral preference for gentle tactile stimulation, suggesting that these afferents contribute to the positive valence of social touch (Liu, Qiao et al. 2022, Liu, Rahman et al. 2025). The ability of brushing to recruit CGRP neurons therefore raises the possibility that CLTMR-driven signals can access ascending circuits that are typically associated with nociceptive and aversive processing. This convergence may reflect a broader role for parabrachial neurons in signaling the salience or motivational significance of somatosensory stimuli, rather than exclusively encoding aversive input. Our data suggest that such affective touch signals are more strongly represented in the parabrachial nucleus than in the SPFp, consistent with the robust activation of lePB neurons observed here.

Both the SPFp and lePB send projections to the amygdala, although they target distinct subregions. CGRP neurons in the SPFp project predominantly to the lateral amygdala, whereas those in the lePB project strongly to the central amygdala. These amygdala nuclei are known to play key roles in threat learning and defensive behaviors. The recruitment of these circuits by non-aversive tactile stimulation may therefore reflect the involvement of these pathways in broader affective or salience processing rather than aversion alone (Kang, Liu et al. 2022, Pauli, Chen et al. 2022). For example, activation of inhibitory neurons within the central amygdala has been shown to suppress pain responses (Hua, Chen et al. 2020), suggesting that a subpopulation of parabrachial inputs may contribute to the modulation of nociceptive processing.

In addition to CGRP neurons in the lePB, our stimuli also activated neurons in the lateral dorsal parabrachial nucleus (ldPB), which contains a population of prodynorphin-expressing neurons that project to targets distinct from those of CGRP neurons (Huang, Grady et al. 2021). Whether the neurons activated in the present study correspond to this population remains to be determined. It is possible that soft tactile stimulation engages parallel parabrachial circuits involving both CGRP and prodynorphin neurons.

Finally, parabrachial circuits may influence sensory processing through descending pathways. Prodynorphin neurons in the ldPB send descending projections to the spinal dorsal horn and have been implicated in the development of mechanical allodynia (Huo, Du et al. 2023). Although the descending projections of CGRP neurons remain less well characterized, parabrachial outputs to hindbrain and brainstem structures may similarly participate in the modulation of sensory processing.

In summary, our findings demonstrate that CGRP neuronal populations in the lePB and SPFp, previously associated primarily with aversive signaling, can also be engaged by non-aversive tactile stimulation. In particular, soft affective touch robustly activated neurons in the lePB, whereas the SPFp showed only limited activation. These results suggest that ascending affective pathways involving CGRP neurons may integrate both nociceptive signals and tactile signals with positive valence, potentially allowing shared circuits to encode the motivational or emotional significance of somatosensory stimuli. Future studies will be required to determine how these circuits contribute to the perception of pleasant and unpleasant touch and how they interact with established pain-processing pathways.

## Acknowledgements

This work was supported by the Swedish Research Council (Vetenskapsrådet; grant number 2021-01109 to LSL), The Wilhelm and Martina Lundgren Science Fund Foundation (grant number 2025-SA-5082 to LSL), and The Swedish Brain Foundation (Hjärnfonden, grant number FO2023-0399 to LSL) and The Mary von Sydow, née Wijk, Donation Fund (stipends to MH and AS; grant nr 2025-343 to FA).

